# Evaluating the diagnostic utility of applying a machine learning algorithm to diffusion tensor MRI measures in individuals with major depressive disorder

**DOI:** 10.1101/061119

**Authors:** David M Schnyer, Peter C. Clasen, Christopher Gonzalez, Christopher G Beevers

## Abstract

Using MRI to diagnose mental disorders has been a long-term goal. Despite this, the vast majority of prior neuroimaging work has been descriptive rather than predictive. The current study applies support vector machine (SVM) learning to MRI measures of brain white matter to classify adults with Major Depressive Disorder (MDD) and healthy controls. In a precisely matched group of individuals with MDD (*n* = 25) and healthy controls (*n* = 25), SVM learning accurately (70%) classified patients and controls across a brain map of white matter fractional anisotropy values (FA). The study revealed three main findings: 1) SVM applied to DTI derived FA maps can accurately classify MDD vs. healthy controls; 2) prediction is strongest when only right hemisphere white matter is examined; and 3) removing FA values from a region identified by univariate contrast as significantly different between MDD and healthy controls does not change the SVM accuracy. These results indicate that SVM learning applied to neuroimaging data can classify the presence versus absence of MDD and that predictive information is distributed across brain networks rather than being highly localized. Finally, MDD group differences revealed through typical univariate contrasts do not necessarily reveal patterns that provide accurate predictive information.

## 1. Introduction

A method for objectively identifying the presence or absence of psychiatric disorders, such as major depressive disorder, is a long standing need in psychiatry (Kapur et al., 2012). One promising approach is to use advances in MRI methods and analytics to derive an objective diagnosis. Although mood disorders have been extensively studied with MRI (Drevets et al., 2008; Lorenzetti et al., 2009), including both structural and functional neuroimaging, few studies have used imaging data to classify MDD. The current study examines whether in vivo diffusion tensor MRI (DTI), a measure of white matter microstructure of the brain, can be used to accurately diagnose major depressive disorder (MDD) (Bracht et al., 2015; Versace et al., 2010). Given the view that depression results from vulnerabilities across interconnected brain networks rather than specific brain nodes (Mayberg, 1997; Wang et al., 2016) (Mulders et al., 2015), approaches that look at the underlying white matter structure that connects these networks could provide important diagnostic utility.

Diffusion tensor imaging (DTI) is a technique that utilizes the ability of MRI to tag water molecules and then wait some period of time to determine the extent to which those molecules are microscopically diffused. By measuring multiple spatial directions, vectors can be generated for each brain voxel to quantify the fiber orientation and integrity of white matter pathways within the cerebral cortex. There are a number of different metrics that can be generated from DTI, but scalar measures are more commonly used in MDD as they can be correlated with disease severity and/or symptoms.

Scalar measures are derived from calculations of one or more of the 3 principle directional vectors of the “diffusion tensor” represented as an ellipsoid. One common metric is fractional anisotropy (FA), which is the extent to which diffusion is characterized as anisotropic, or highly directional (high FA) vs unrestricted or isotropic (low FA). For example, one of the white matter pathways with the highest FA values is the corpus callosum, due to its highly organized, densely packed fibers that run mainly in a left-right direction. In addition to directionality, FA is influenced by axon size and density, pathway geometry, and extent of fiber intersections (Alexander et al., 2007; Beaulieu, 2002).

Another scalar measure is calculated as the average of the 3 directional vectors and is referred to as mean diffusivity (MD). MD reflects the extent to which there is water movement at all and is a useful clinical measure to indicate edema and restricted liquid flow. Axial diffusivity (AD) is the strength of the primary directional vector and radial diffusivity (RD) is the mean of the 2 non-principle vectors. While all these measures can be calculated from DTI imaging, FA is the most reliably sensitive measure of between group microstructural white matter differences (see (Feldman et al., 2010).

A number of studies have demonstrated differences in FA values between patients with MDD and healthy controls. A meta-analysis of 11 studies that examined FA in individuals with MDD (Liao et al., 2013) identified 4 consistent locations associated with altered FA in MDD compared to healthy controls: right and left dorsal frontal regions, a region of the right fusiform and a region of the right occipital lobe. A review paper that focused on 35 studies of WM alterations in pathways associated with the reward circuit (Bracht et al., 2015), found reduced FA in the cingulum bundle, increases and decreases of FA in the uncinate fasciculus in adolescents, and reduced FA in the uncinate fasciculus and the anterior thalamic radiation/supero-lateral medial forebrain bundle during acute depressive episodes in adults. Other studies have focused on WM microstructure in those at risk for MDD either by virtue of family history (Keedwell et al., 2012) or genetic polymorphisms (Pacheco et al., 2009).

Given the heterogeneity of findings, an important theme that emerges from this work is that white matter microstructure alterations in MDD are distributed across many defined brain networks. Thus, the use of DTI to understand underlying WM features associated with MDD has been useful in characterizing the underlying brain circuits associated with the psychiatric disorder. However, despite these interesting results, it remains unclear if and when DTI might be implemented as a promising diagnostic tool. One of the steps needed in order to accomplish this goal would be to quantitatively determine how well DTI measures can discriminate people with and without MDD.

One approach to examine the diagnostic utility of MRI modalities involves applying multivariate machine learning classification algorithms in order to identify individuals with a specific disorder (Orrù et al., 2012). There has been increasing interest in applying multivariate pattern analysis methods in order to categorize patients suffering from psychiatric disorders from healthy controls (Cohen et al., 2011). The main advantage of these approaches is that they are predictive. Once a classifier has been defined, it can then be tested on new individuals to predict group membership. These approaches have utilized functional brain imaging (Craddock et al., 2009; Zeng et al., 2012) and structural brain images (Ardekani et al., 2011). This approach is starting to be applied to MDD (for review see (Patel et al., 2016)).

To date, this machine learning approach has been applied to a range of MRI modalities in an effort to automate the diagnosis of a number of disorders (Magnin et al., 2009) but few studies have been completed with MDD. One of the earliest studies examined the application of SVM to depression diagnosis using resting-state fMRI in 20 patients with MDD and 20 matched controls. While the purpose of this work was to examine different feature selection approaches, the unselected data yielded only modest classification accuracy – 62.5%. In another study of 32 women (14 with MDD), researchers applied global tractography-based graph metrics for the classification of depression (Sacchet, 2015). The investigators characterized connectivity between 34 cortical regions resulting in 9 global graph metrics that were then used in a SVM classification. Combined, the 9 metrics classified MDD and controls at a performance level of 71.9% accuracy.

A second study applying SVM classification to DTI in order to study depression applied probabilistic tractography to reconstruct specific WM tracts and then extracted anatomical networks (Fang et al., 2012). SVM was then applied to determine the most discriminating connections within these networks. The resulting classifications were highly accurate (91.7%) and revealed that the most discriminating connections were primarily within the cortical-limbic network where it was revealed that young adult first episode MDD patients displayed increased anatomical connectivity relative to healthy controls. In this study, a two sample t-test approach was taken to select features to be utilized in classification. An important limitation of the use of feature selection algorithms can often produce sample-specific results that may not generalize to new data.

The aim of the current study is to continue to explore the utility of DTI in the classification of individuals diagnosed with MDD. The previous examinations using SVM classification of DTI imaging in MDD did not report the utility of standard scalar metrics such as FA, MD, RD and therefore, those metrics will be examined here. Moreover, when feature selection techniques were applied (Fang et al., 2012), classification accuracy was greatly increased. It is important to examine the predictive power of classification both with and without feature selection to understand the predictive range of these techniques. This approach was applied to a sample of treatment-seeking participants with DSM-IV Major Depressive Disorder who were part of a study testing the efficacy of attention bias modification (Beevers et al., 2015).

## 2. Methods

### 2.1 Sample

Fifty-two treatment-seeking participants with DSM-IV Major Depressive Disorder (MDD) and 45 healthy control (HC) participants were recruited for this study from advertisements placed online, in newspapers, and on late-night TV. Participants were screened for medical or physical conditions that would preclude participation in an fMRI study (e.g., orthodontic braces). They also completed an abbreviated Mini International Neuropsychiatric Interview (MINI) (Sheehan et al., 1998) to determine provisional MDD diagnosis (MDD) or absence of psychiatric symptoms (HC). Diagnoses were subsequently confirmed in-person with a Structured Clinical Interview for the DSM-IV Disorders (SCID) administered by a trained research assistant.

Participants in the MDD group met diagnostic criteria for current major depressive disorder, but did not meet criteria for substance abuse (past year) or dependence, current or past psychotic disorder, bipolar disorder, and/or schizophrenia. Participants in the HC group did not meet criteria for any current or past psychiatric disorder. Consistent with previous research (Amir et al., 2009; Sheehan et al., 1998), participants receiving pharmacological treatment were allowed into the study if there had been no medication change in the 12 weeks prior to study entry.

In order to increase the likelihood that groups were matched for structural brain characteristics, we selected from the larger study sample a subset of healthy control participants that were matched for age and gender on a 1-to-1 basis with individuals from the MDD group. To minimize brain changes associated with aging, selected participants were also between 18 and 35 years of age. This matching algorithm resulted in a sample size of 50 participants (25 MDD, 25 healthy controls). This sample was used for all analyses reported below.

### 2.2 Imaging Methods

#### 2.2.1 Acquisition

All scanning was performed on a whole body 3T GE MRI scanner (Excite) with an 8-channel head coil. The primary measure of white matter (WM) was derived from a HARDI diffusion MRI that was collected using single shot echo planar imaging, and a twice-refocused spin echo pulse sequence, optimized to minimize eddy current-induced distortions (GE 3T, TR/TE=12000/71.1, B=1000, 128-by-128 matrix, 3mm (0-mm gap) slice thickness, 1 T2 + 25 DWI). Forty-one slices were acquired in the approximate AC-PC plane. The 25 diffusion weighted directions resulted in a high signal-to-noise diffusion volume that took approximately 7 minutes to acquire. Participant head motion was minimized by instruction and the use of foam inserts.

#### 2.2.2 Diffusion Tensor Processing

All diffusion image analysis was conducted with the FMRIB Software Library (FSL, http://www.fmrib.ox.ac.uk/fsl). First, images were corrected for eddy current distortions and for motion using the b=0 volume as a reference. The registered images were skull-stripped using BET. Diffusion tensors were then calculated on a voxel by voxel basis using conventional reconstruction methods (Basser et al., 1994) and from these tensors Fractional Anisotropy (FA), Mean Diffusivity (MD), Axial Diffusivity (AD) and Radial Diffusivity (RD) maps were calculated on a voxel-by-voxel basis.

Individual FA maps were then entered into the TBSS (Tract-Based Spatial Statistics) pipeline (TBSS; Smith et al., 2006; 2009). The critical steps of the TBSS pipeline include: (1) FA data were aligned onto the common FMRIB58 FA template (MNI152 standard space) using a non-linear registration algorithm FNIRT; (2) a mean FA image was created for each group (MDD and NC separately) from the images for all the subjects in MNI152 space; (3) images are then thinned to generate a mean FA white matter skeleton that represented the center of all tracts common to the entire group; (4) the resulting map was thresholded to FA values greater than 0.2 in order to exclude gray matter and low intensity voxels that may reflect partial volume effects with gray matter; (5) the aligned FA volume for each subject was then projected onto the skeleton by filling the skeleton with FA values from the center of the nearest tract; (6) this is achieved for each skeleton voxel by searching perpendicular to the local skeleton structure for the maximum value in the FA image of the subject; (7) for the purposes of this analysis, FA values were used from each subject for both the total FA map as well as the FA skeleton map; (8) the remaining scalar measures (MD, AD, and RD) were processed using the alignment parameters from the FA processing stream in order to generate common space maps that are true to the white matter architecture.

### 2.31 TBSS Voxel-wise Analysis

In order to compare results from univariate contrasts of WM maps to those obtained through SVM classification voxel-wise comparisons were conducted between the final group of 25 MDD patients and 25 healthy controls. Nonparametric statistical comparisons were performed on the skeletonized FA images using the FMRIB Software Library (FSL) randomize algorithm based on permutation generated statistical thresholds, with corrections for multiple voxel-wise comparisons using threshold-free cluster enhancement (TFCE). Anatomic locations of voxel clusters with statistically significant differences in FA between MDD and HC at *p* < 0.05 were determined using 1000 permutations. The same approach was taken to examine the MD, AD and RD maps.

### 2.3.2 Support Vector Machine Classification Analysis

Classification of individual subjects was undertaken using the freely available Pattern Recognition for Neuroimaging Toolbox (PRoNTo-http://www.mlnl.cs.ucl.ac.uk/pronto/ (Schrouff et al., 2013)). The Linear Support Vector Machine (SVM) is conceptually illustrated in Figure 1. Each dimension corresponds to a feature set (scalar voxel values reduced through the kernel process in this case) and thus each subject is located in the space depending upon its constituent features. The pluses and minuses constitute the two putative categories, namely MDD and HCs. The SVM finds what is known as the maximum margin decision boundary, which is the hyperplane that is furthest from the least discriminating features of the to be discriminated categories. The hyperplane is also associated with a maximum margin that best separates the two groups, where larger margins are associated with better classifier generalizability. The margin is fully specified by the subset of training samples that lie on it and reflect the support vectors, since they represent the specific cases that support the solution (See Figure 1).

**Figure 1.**
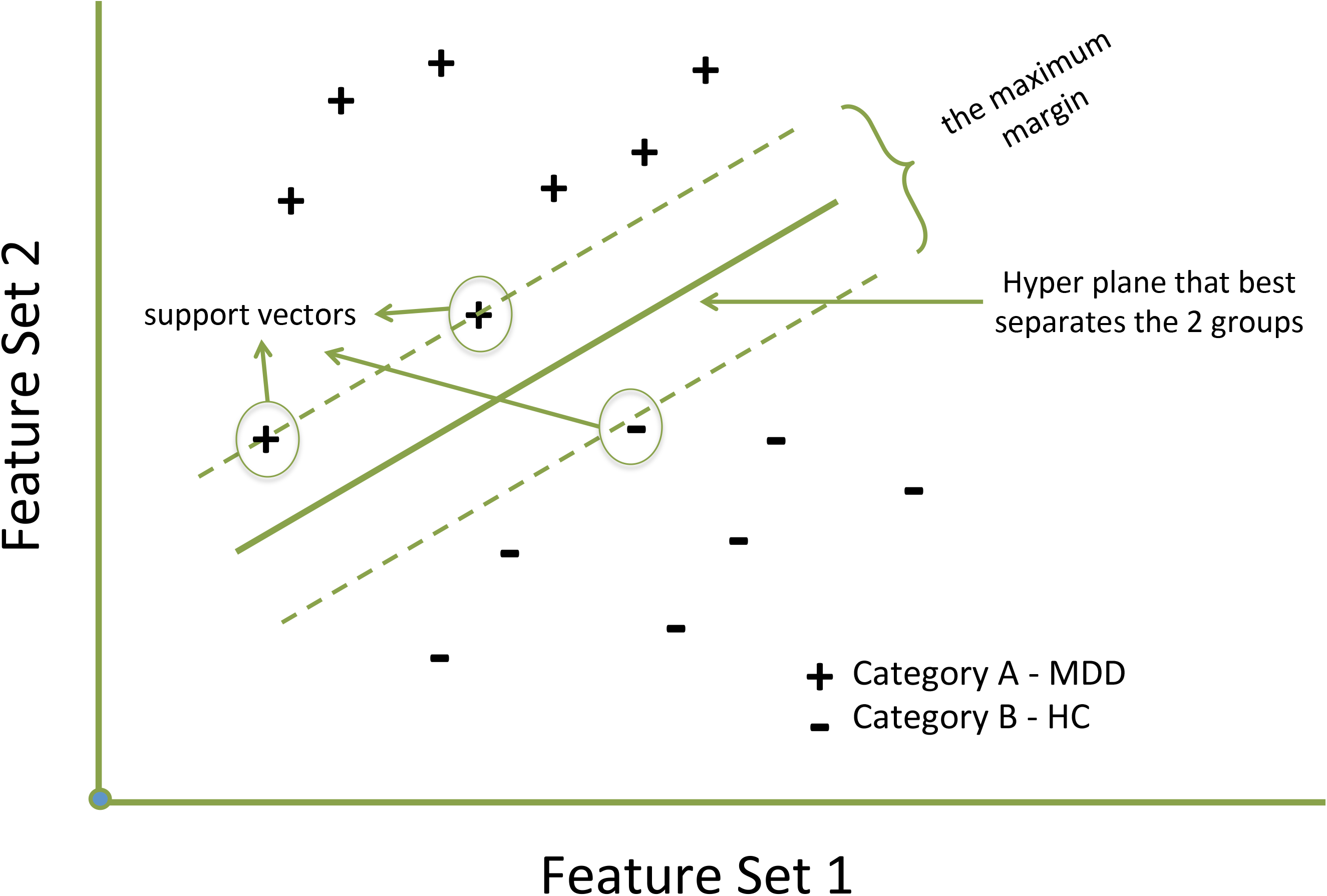
A hypothetical graphic of application of support vector machine algorithms in order to classify 2 categories. Two feature sets can be plotted against one another and a hyperplane generated that best separates the groups based on the selected features. The maximum margin represents the margin that maximizes the divide between groups. Cases that lie on this maximum margin define the support vectors.

The SVM approach in PRoNTo utilizes LIBSVM for matlab, which is an implementation of a linear-kernel SVM for binary classification (Chang and Lin, 2011).

Following DTI analysis, the resulting scalar maps were prepared for classification using 3 different approaches. First, whole brain scalar maps were used, although masked to include only white matter by thresholding the group mean FA map at a value of FA > 0.2. The resulting image was then used as a mask to select voxels for each participant as input to the classification analysis. Using this approach, classification was conducted on the resulting whole brain maps, a right hemisphere map and a left hemisphere map. This same approach was taken to examine the FA skeleton map, namely classification was conducted on the whole brain skeleton and a left and right hemisphere only skeleton. Finally, a “feature selection” approach was taken only for the FA map^1^ in order to reduce the number of voxels to a subset most relevant for classification (Mwangi et al., 2013).

This approach is a feature-wise *t*-test filter (TF) to determine features that have different group means (Mitchell et al., 2004). This step eliminates non-discriminative voxels that reduce classification accuracy. This was accomplished by first removing one pair of MDD and HC participants from the dataset and then splitting the 2 groups of remaining participants in half randomly and contrasting MDDs vs HCs in each split half data separately. Resulting t-maps were thresholded at *t* = 2.10 (95% confidence interval) and then the 2 split half t-maps were combined to retain all thresholded voxels in common between them (*p* < 0.0025). SVM classification was then applied to this overlap map and accuracy was tested for each map utilizing the participant pair left out.

This procedure produced 25 separate TF maps and the average accuracy scores were calculated from the 25 testing folds. This average number of voxels across the 25 separate TF maps was 1232mm^3^ voxels with a standard deviation of 487mm^3^ voxels. In summary, 8 datasets were created; the FA > 0.2 thresholded FA, MD, AD and RD maps for the whole brain, and the right and left hemispheres, the whole brain FA skeleton map and skeleton right and left hemispheres, and finally FA maps that were feature selected using the split-half *t*-test filtering technique.

Following data-selection, classification was then carried out using the Support Vector Machine (SVM) approach. For more detail of the processing steps the reader is referred to Schrouff et al. (for flow chart of processing steps see Figure 1 in Schrouff et al., 2013). The first step is the generation of a “similarity matrix” in the form of a linear kernel (Hofmann et al., 2008) that reduces the dimensionality of the input data set to a matrix the size of N_samples_ by N_samples_. This kernel matrix is a similarity measure resulting from the dot product of all brain voxels reflecting their degree of similarity. This matrix is then input into the classification algorithm. For classification, two classes were defined – MDD and HC and processed using a soft-margin hyper-parameter approach. In order to examine the model’s estimation power, a leave-one-subject-per-group-out (LOSPGO) cross-validation approach was used. In each step of the cross-validation, the individuals are grouped into disjoint training and testing sets such that there are no subjects used for both training and testing in a single step. This process is repeated across all pairs of left out individuals and the results from each step are averaged to obtain a final estimate of classification accuracy. The model performance was tested for significance using permutation testing where the model was estimated 1000 times with randomly permuted class labels that produces a *p*-value for each of the performance values.

In order to assign classification power to specific locations in the brain WM, PRoNTo takes the linear SVM models and recovers model weights and transforms the weights vector into a map in the original image (voxel) space. These maps contain at each voxel the corresponding weight of the linear model that reflects how much this particular voxel contributed to the classification. In addition, the contribution of specific regions within the ICBM-DTI-81 white-matter atlas (Mori et al., 2008), an atlas where 48 white matter tract labels were created by hand segmentation of a standard-space average of diffusion MRI tensor maps from 81 subjects. This is accomplished by first summing the absolute values of the weights within each region divided by the number of voxels in that region. Then the contribution of each region is divided by the total contribution of all regions resulting in values that reflect the percent contribution of each region to the decision function. These regions can then be ranked by descending order based on their contribution to the model and examined to understand how regions contribute to the classification accuracy.

## 3. Results

### 3.1 Demographic Characteristics

Table 1 shows the demographic and depression symptom profile of the MDD and HC groups. The groups were well matched on age, gender and income but were marginally different on ethnic distribution. Given the MDD diagnosis, the groups were significantly different on BDI-II and IDAS (Inventory of Depression and Anxiety Symptoms, (Watson et al., 2008)).

**Table 1.** Table of participant demographics. MDD and HC significantly differed only on depressive symptoms.

|  |  | MDD | HC | Test<br>Statistic | *p*-value |
| --- | --- | --- | --- | --- | --- |
| Age | | 23.4 (3.6) | 23.5 (4.0) | $t = 0.12$ | 0.91 |
|  | Age Range | 18-31 | 19-33 |  |  |
| Gender | | 12 female | 12 female | $X = 0$ | 1 |
| Ethnicity | | 17 Caucasian | 11 Caucasian | $X = 2.92$ | 0.09 |
| Income | | 46614 (36K) | 54640 (25K) | $t = 0.88$ | 0.38 |
|  | Income Range | 5K-150K | 12K-100K |  |  |
| BDI | | 35.1 (8.1) | 2.4 (3.0) | $t = 18.86$ | < 0.001 |
|  | BDI range | 21-47 | 0-12 |  |  |
| IDAS | | 74.0 (8.8) | 28.8 (5.0) | $t = 22.50$ | < 0.001 |
|  | IDAS range | 61-90 | 22-38 |  |  |
| BDI-II - Beck Depression Inventory-II |  |  |  |  |  |
| IDAS - Inventory of Depression and Anxiety Symptoms (Watson et al., 2008). |  |  |  |  |  |

### 3.2 TBSS voxel-wise results

Contrasting the FA skeleton between MDD and HC groups revealed only a single significant cluster: FA values were greater for MDD than HC in the right body of corpus callosum (see Figure 2). Contrasting the AD skeleton between MDD and HC groups revealed a single significant cluster: lower AD values for MDD vs HC in the right anterior thalamic radiation (see Figure 3). The skeletonized MD map revealed a very similar effect of lower MD in the anterior thalamic radiation in MDD versus HC. In this case, the effect was bilateral. Finally, the RD skeleton map did not reveal any significant depression group differences using the univariate voxel-wise contrast approach with cluster wise corrections.

**Figure 2.**
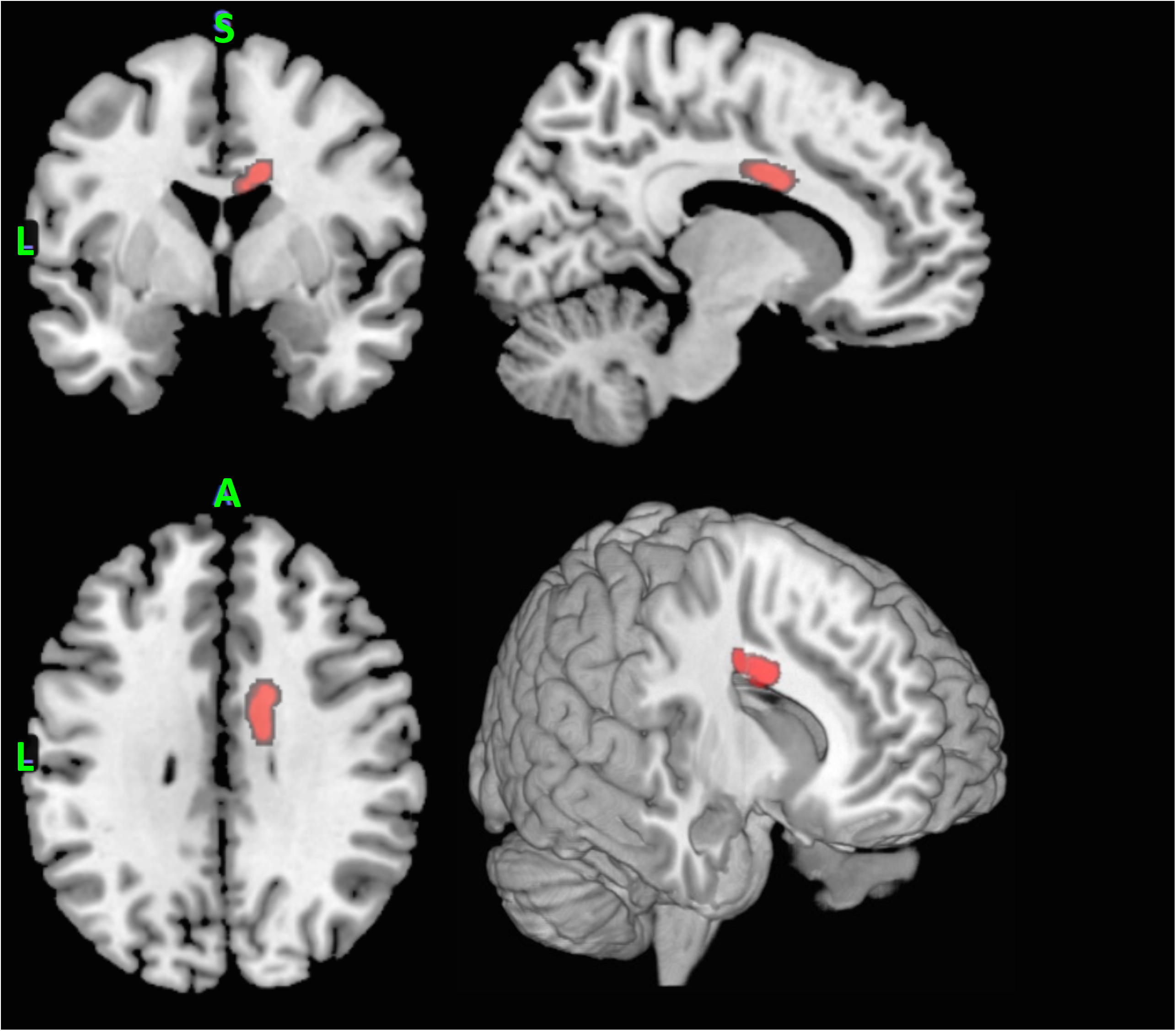
Region of right body of the corpus callosum that revealed significantly higher FA values in MDD relative to HCs. From the top left clockwise – coronal, sagittal, 3D render and axial projections.

**Figure 3.**
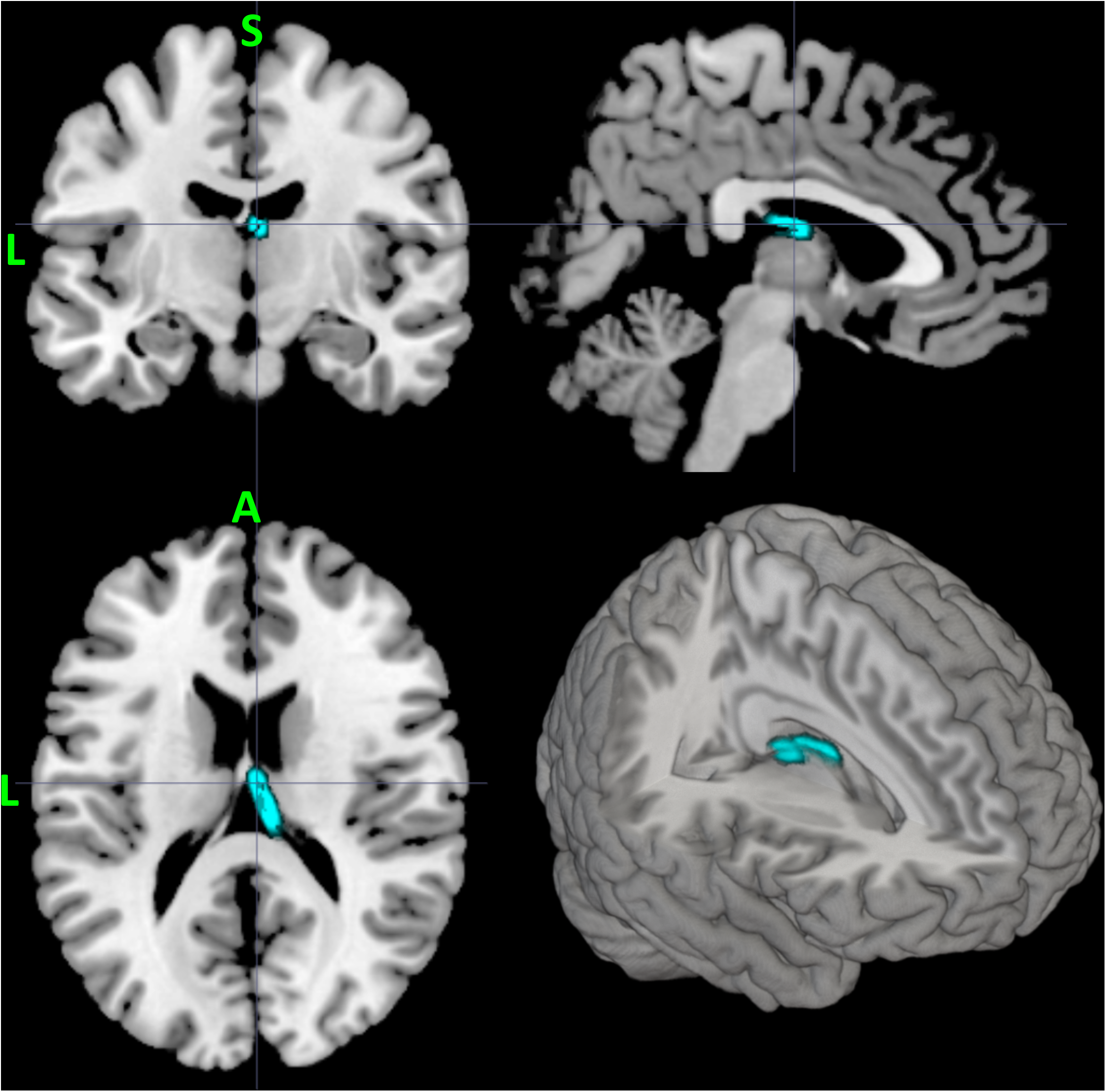
Region of anterior thalamic radiation that revealed significantly lower AD and MD values in MDD relative to HCs. From the top left clockwise – coronal, sagittal, 3D render and axial projections.

### 3.3 Support vector machine classification results

Accuracy is the total number of correctly classified test samples from each leave-one-pair-out set divided by the total number of test samples, irrespective of class. Classification assessment across the 8 datasets indicated that all sets classified MDDs significantly (see Table 2). For the whole brain FA map total classification accuracy was 70.0% (permutation *p* = 0.015) with a specificity of 80.0% and a sensitivity of 60.0%. This performance was improved when just the right hemisphere FA map was used, resulting in total accuracy of 74.0% (permutation *p* = 0.005), with a corresponding specificity of 80.0% and sensitivity of 68.0%. The whole brain skeleton map was comparable to whole brain FA, as total classification accuracy was 70.0% (permutation *p* = .011) with a specificity of 84.0% and a sensitivity of 56.0%. Testing the hemispheres separately did not improve performance of the FA skeleton maps.

**Table 2.** Table of SVM classification performance across the 8 different fractional anisotropy (FA) mapping profiles. Values in parenthesis indicate resulting *p* value from permutation testing. Given that only a single fold was tested for each of the 25 TF selected maps, a *p* value could not be calculated.

| MAP | Voxels<br>2mm<br>cubic | Classification Accuracy<br>(permutation p values) |  |  |
| --- | --- | --- | --- | --- |
|  |  | Total | MDD | HC |
| **Whole Brain FA** |  |  |  |  |
| All FA | 1,083,793 | 70 (0.015) | 80 (0.003) | 60 (0.23) |
| Right Hemisphere FA | 168,929 | 74 (0.005) | 80 (0.004) | 68 (0.06) |
| Right Hemisphere FA -W/O CC | 165,035 | 76 (0.002) | 84 (0.001) | 68 (0.07) |
| Left Hemisphere FA | 164,399 | 68 (0.03) | 72 (0.022) | 64 (0.122) |
| **Whole Brain FA Skeleton** |  |  |  |  |
| All FA Skeleton | 111,052 | 70 (0.011) | 84 (0.001) | 56 (0.363) |
| Right Hemisphere FA Skeleton | 75,939 | 68 (0.020) | 72 (0.017) | 64 (0.108) |
| Left Hemisphere FA Skeleton | 75,804 | 68 (0.021) | 76 (0.005) | 60 (0.219) |
| **Feature Selected** |  |  |  |  |
| TF Split-Half Selected FA | 1231<br>(SD 487) | 74 | 68 | 76 |

Results for the MD, AD and RD maps were less promising (see Table 3). For MD, the whole brain significantly classified MDD but not healthy controls. The left and right MD maps were unsuccessful at significantly classifying either group. The AD maps showed a similar pattern to the MD map, namely only the whole brain map significantly classified MDD but not HC. Finally, the RD maps were not able to significantly classify any group. Because of the failure of these 3 scalar measures to significantly distinguish between groups, a feature selection approach was not taken. Finally, the t-test filtered feature selected FA map did not increase the performance of the SVM above what was seen in with the whole brain right hemisphere FA map. Total classification accuracy was 74.0% with a specificity of 68.0% and a sensitivity of 76.0%.

**Table 3.** Table of SVM classification performance across the 3 different mapping profiles for the scalar values of MD, AD and RD. Values in parenthesis indicate resulting *p* value from permutation testing.

| MAP | Classification Accuracy<br>(permutation $p$ values) | | |
| --- | --- | --- | --- |
|  | Total | MDD | HC |
| **Whole Brain MD** |  |  |  |
| All MD | 70 (0.002) | 96 (0.057) | 44 (0.517) |
| Right Hemisphere MD | 68 (.006) | 88 (0.155) | 48 (0.499) |
| Left Hemispher MD | 58 (.095) | 92 (0.132) | 24 (0.693) |
| **Whole Brain AD** |  |  |  |
| All AD | 74 (.001) | 96 (0.048) | 52 (0.472) |
| Right Hemisphere AD | 72 (.001) | 88 (0.089) | 56 (0.447) |
| Left Hemisphere AD | 62 (.056) | 88 (0.141) | 36 (0.620) |
| **Whole Brain RD** |  |  |  |
| All RD | 58 (.164) | 80 (0.350) | 36 (0.499) |
| Right Hemisphere RD | 52 (.609) | 84 (0.341) | 20 (0.827) |
| Left Hemisphere RD | 52 (.612) | 84 (0.324) | 20 (0.831) |

**Table 4.**
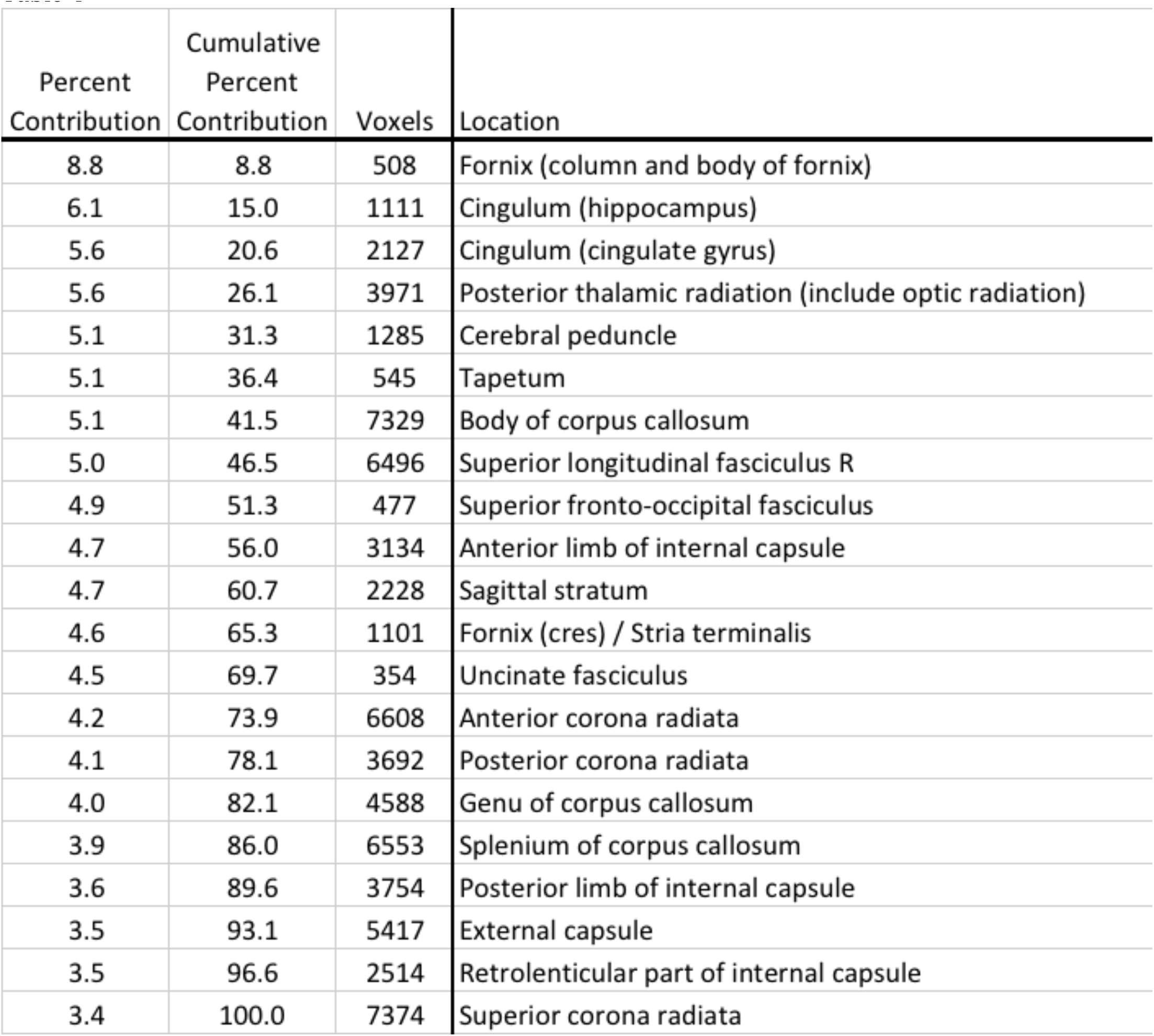
Table of weight map values generated for the right hemisphere FA map defined in terms of ICBM-DTI-81 atlas regions. Includes the percent contribution and total voxels for each region.

A prediction plot for the best performing dataset, namely the right hemisphere whole-brain FA map can be seen in Figure 4. The plot displays the output “decision function values” where positive numbers represent the MDD class and negative numbers the HCs. The zero line is the decision bound for this classifier. A well-performing classifier will show clear separation of the 2 classes. In addition, there appears to be a wider range of variability in the decision function values for HC when compared to MDD.

**Figure 4.**
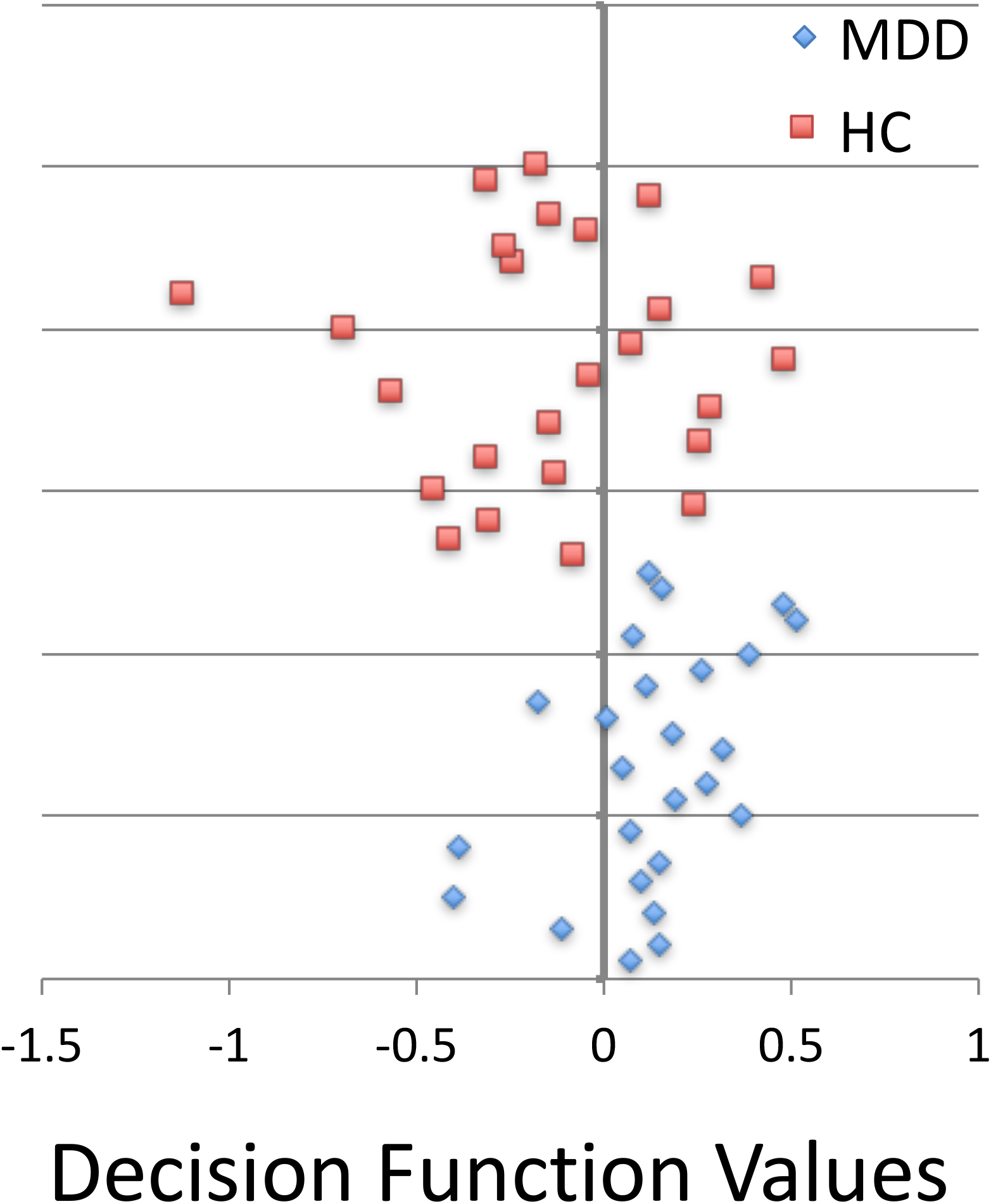
Results of the leave-one-pair-out test of SVM accuracy for the right-hemisphere FA map. Normalized decision function values are plotted for MDD (blue triangles) and HC (red squares) participants. The zero line represents the decision boundary.

Given that neuroimaging data contains spatial information that may be critical in understanding the underlying WM pathways that contribute to classification accuracy, PRoNTo allows one to generate a “weight map”. The weight map is a spatial representation of the decision function where each voxel contributes with a certain weight to the classifier decision function. A weight map was generated for the right hemisphere FA map using the ICBM-DTI-81 atlas. The results of this map, defined in terms of ICBM-DTI -81 ROIs, can be seen in Table 3 and the projections of these ROIs on a standard brain can be seen in Figure 5. The table includes regions ranked by their total contribution to the model in descending order, the cumulative percent contribution, the number of voxels within that region and finally the ROI WM label.

**Figure 5.**
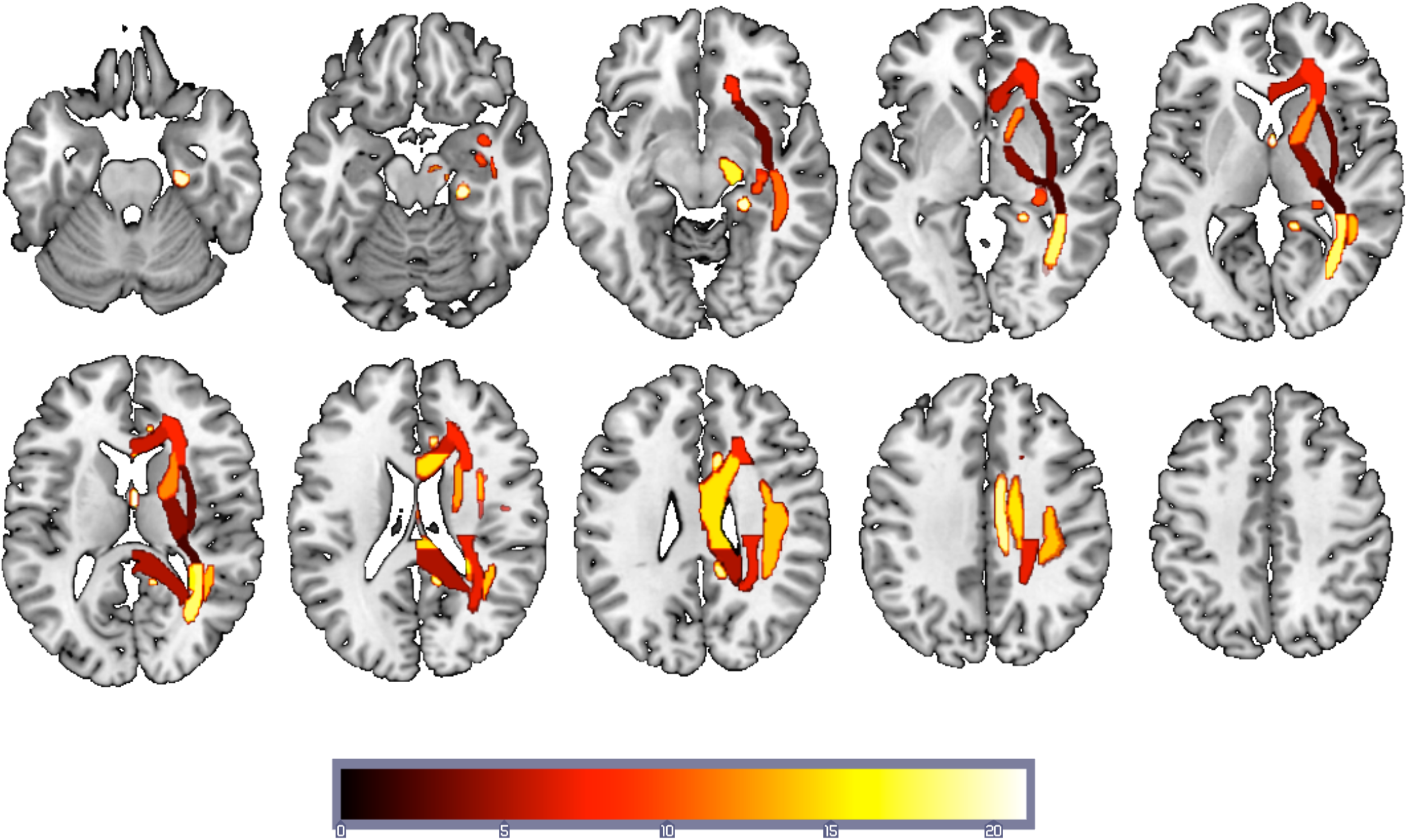
Regions supporting the SVM solution for the right-hemisphere FA map masked using ICBM-DTI-81 white-matter atlas map projected onto a standard T1 weighted image. The heat color map reflects the rank ordering of the contribution to the SVM with hot being the region of greatest contribution (Fornix - column and body of fornix).

The second largest contributor to the model, in terms of number of voxels, lay in the right body of the corpus callosum and overlaps with the region revealed in the TBSS univariate analysis. However, this region makes a relatively small contribution to the classification—an estimated 5.1%. In order to confirm the contribution of this region to classification accuracy was minimal, a nested model test was performed by removing the voxels contained within this region and then the classification was rerun. Removal of these 7329 voxels resulted in numerically higher classification accuracy - total classification accuracy of 76.0% with a specificity of 84.0% and a sensitivity of 68.0%, although likely this change was not significant.

## 4. Discussion

The current report supports using machine learning algorithms to capture the diagnostic information contained in structural MRI data in order to differentiate between patients diagnosed with MDD and healthy controls. Despite the relatively small sample size, using both an unselected and a feature selected DTI dataset, support vector machine binary classification was able to significantly distinguish between MDD and HC using the DTI metric of fractional anisotropy (FA). While this approach awaits demonstrated predictive power when applied to an independent dataset, nevertheless there is important information that can be derived from this project.

Across multiple DTI metrics – FA, MD and AD there were several findings from the univariate between group contrasts that are informative and consistent with previous reports in patients with MDD. First, examining the FA skeleton revealed that a region of the right body of the corpus colosum (CC) had higher FA values in MDD relative to HC. While some reports have demonstrated that depression is associated with decreased FA values in the body of the CC (Cole et al., 2012), a study by (Frodl et al., 2012) found that unaffected first-degree healthy relatives (UHRs) of patients with MDD revealed increased FA values in the CC, which they speculated might represent vulnerable characteristics for the formation of depression. Finally, a meta-analysis of WM abnormalities in MDD that examined 17 studies that included 641 MDD patients and 581 HC (Chen et al., 2016) found that the CC consistently reveals differences between MDD and HC, particularly in the genu and body regions, even if the directionality of those differences remains unclear.

The region of CC revealed in the current study is right lateralized and clearly involves a region of interface with the superior longitudinal fasciculus (SLF). Greater FA in this region may indicate a reduction in fiber complexity (Beaulieu, 2002) in a location where one would expect increased crossing fibers between the CC and SLF. It is important to be careful when interpreting the directionality of FA differences between groups since FA is sensitive to different elements of microstructure depending on location (Beaulieu, 2002). Moreover, it is not always clear whether investigators tested the reverse contrast of MDD > HC as many articles do not explicitly state whether any regions showed greater FA in MDD relative to HC. Finally, the right body of the CC lies along the midline of the brain corresponding to a functional network referred to as the default mode network (DMN). Greater functional connectivity within the DMN has been consistently associated with MDD (Hamilton et al., 2015). Increased FA in the body of the CC may be a structural correlate of increased functional connectivity in the DMN observed in MDD.

In addition to the finding of higher FA values in the right CC, findings in the anterior thalamic radiation indicated lower axial diffusivity and lower mean diffusivity in MDD relative to HC. This is consistent with a number of previous studies (Lai and Wu, 2014) (Korgaonkar et al., 2014) that have shown alterations in the anterior thalamic radiation associated with MDD. In a whole-brain examination of WM structural networks in MDD relative to controls, differences were revealed in two brain networks (Korgaonkar et al., 2014), one being a frontal-subcortical network that included regions of frontal cortex, the caudate and the thalamus. In the current work, the anterior thalamic radiation revealed decreases in the longitudinal component, possibly reflecting reduced myelination in MDD patients and therefore lower connectivity between the thalamus and the cortex.

While the univariate results add information to the corpus of work examining what brain structural features are associated with depression, it is inherently an explanatory approach to the data (Yarkoni and Westfall, 2016). By contrast, the results of our machine learning classification hold the promise of predicting who is depressed or even potentially who might be vulnerable to depression.

Across the four scalar measures derived from the DTI imaging, only the results from the FA maps provided significant prediction accuracy. Using a whole brain approach, the right hemisphere selected FA map predicted MDD patients with 80% accuracy and healthy controls with 68% accuracy (total accuracy was 75%). Restricting the SVM learning to just the skeletal FA values resulted in lower prediction accuracy overall, indicating that restricting the map to the highest FA path through the brain does not increase predictive ability. Finally, using a TF feature selection approach using the voxels which most clearly separated the groups did not improve prediction accuracy over whole-brain FA.

It is interesting to note that of the four metrics examined, FA was uniquely successful at classification. One possible explanation for this finding is that FA reflects a composite of the other scalar measures and thus maximizes differences that might be distributed across the other measures. This is consistent with considerable research showing FA to be a highly sensitive but fairly nonspecific measure of white matter microstructure and white matter neuropathology (Alexander et al., 2007).

A number of aspects of the SVM results are worth exploring. First, there was a clear difference in the predictive power of right hemisphere FA values relative to left. In the unselected data set it was the right hemisphere FA maps that clearly classified both MDD and HCs. This finding is consistent with work showing that individual differences in the right hemisphere may be critically tied to depression (Bruder et al., 2012; Costafreda et al., 2009; Talarowska et al., 2011) and depression vulnerability (Beevers et al., 2010; Clasen et al., 2013). For instance, a meta-analysis of 10 whole-brain-based FDG-PET studies in MDD revealed decreased cerebral metabolism in the right caudate, right insula and right cingulate gyrus (Su et al., 2014) and an older study found in a small group of moderate to severely depressed patients, reduced right hemisphere metabolism in the superior temporal lobe (Post et al., 1987). The current study examines the predictive ability of WM measures across the whole brain and the results point to the importance of the right hemisphere in that endeavor.

Another interesting aspect of the current work is the finding that a region revealed in the univariate examination of between group differences, the right body of the CC, does not appear to play a significant role in SVM classification accuracy. Taking a univariate approach to examining the WM differences between groups assumes that the spatial units of the WM are independent entities. As such, each individual image element is tested separately and does not take into account distributed relationships between elements (McIntosh and Mišić, 2013). The univariate finding could easily reflect a subset of participants for whom there are particularly large differences in FA values between MDD and controls. However, the resulting statistical difference is clearly not helpful in determining which group any given individual belongs to. This finding is very important to underscore since the vast majority of neuroimaging correlates of depression are never tested for their predictive value.

An early demonstration of the utility of multivariate approaches to neuroimaging revealed that removing peak fMRI activation differences between conditions did not significantly change the performance of a classifier trained across a distributed network (Haxby et al., 2001). This work was one of the first to reveal the utility of examining distributed elements in order to best discriminate between conditions (Norman et al., 2006). Therefore, the SVM analysis, which is fundamentally a multivariate approach, can capture a distributed network of elements that contribute to the summed ability to separate MDD patients from HCs. The greater the distribution of spatial elements the more unique the solution can become. This perspective is supported by a recent paper where it was shown that “high value” hubs of human brain networks are more likely to be anatomically abnormal across multiple brain disorders including depression (Crossley et al., 2014). This approach does have its drawbacks in that the best solution to separate the groups may not generalize well outside of a specific dataset. For this reason, accuracy was tested in the current studying using a leave-one-pair-out approach but it will fall on future work to examine the performance of trained classifiers on an independent dataset. For the time being, these results help set some boundaries for what might be expected for the predictive power of DTI with respect to MDD.

While the distributed nature of the brain regions associated with accurate classification of MDD patients is a strength when applied to diagnostics, it is also a challenge when it comes to understanding the contribution of specific brain nodes to the disorder. White matter FA values in elements across all three brain networks previously associated with MDD (Mulders et al., 2015) – the DMN, the salience network (SN) and the central executive network (CEN) – also known as the frontal-parietal network (FPN) contributed to the accuracy of classification (see Figure 4). Therefore, it is clear from the current study that the most accurate prediction of MDD from white matter microstructure is obtained when including distributed networks across the brain. It follows that it is likely not possible to derive a highly accurate MDD classification using one white matter tract or network. Fortunately, advances in machine learning and related statistical techniques allow for the integration of highly dimensional data into prediction algorithms. These methods therefore appear to have substantial promise for the development of diagnostic tools that can objectively classify the presence or absence of major depressive disorder and other psychiatric disorders.

## Acknowledgements

This research was supported in part by an award from the National Institute of Health (R21MH092430). The content is solely the responsibility of the authors and does not necessarily represent the official views of the National Institutes of Health. Special thanks to Robert Chapman for his help with data collection. We would also like to thank Ian Dobbins for his assistance with the support vector machine figure. P.C. Clasen is now affiliated with Facebook.

## Footnotes

1 A feature selection approach was only taken with the FA measure since none of the other measures resulted in any significant univariate contrast results, nor SVM significant classifications.

